# The development of the brain was revealed by brain entropy (BEN) during movie watching from the age of three to twelve years

**DOI:** 10.1101/2025.10.19.683278

**Authors:** Donghui Song

## Abstract

Mapping the developmental landscape of brain function is a key objective in modern neuroscience, greatly advanced by the public release of neuroimaging datasets. Here, we utilized movie-watching fMRI to study brain entropy (BEN) in 3-12-year-olds from OpenNeuro. The results of the analysis revealed that decreasing BEN within the action-mode network (AMN) serves as a key developmental marker in children. The AMN underpins goal-directed cognition, and its entropy reduction may signify a crucial shift from early, externally driven sensorimotor processing toward the emergence of more internalized, self-referential thought processes associated with the default-mode network. These findings highlight BEN as a powerful and sensitive tool for deciphering the principles of functional brain development.

## 1. Introduction

Understanding the developmental mechanisms of brain function represents a fundamental pursuit in neuroscience. The advent and public sharing of large-scale neuroimaging datasets have been instrumental in propelling this field forward, establishing the mapping of lifespan brain development as a key research direction. Early work in this area focused on creating comprehensive charts of age-related changes in brain morphology. Building on this foundation, subsequent research has expanded the scope to investigate the dynamic development of functional and structural brain networks throughout life. Together, these converging lines of inquiry provide a vital macroscopic perspective, greatly advancing our understanding of typical brain development and offering a framework for identifying general principles that govern the maturing brain.

Despite these advances, the transition from mapping brain morphology (Bethlehem et al., 2022) to brain network (Sun et al., 2025) reveals a significant knowledge gap, and the question of how local brain function changes throughout development remains unanswered. Here, brain entropy (BEN) (Wang, Li, Childress, & Detre, 2014) emerges as a pivotal measure. As a metric that sensitively captures the dynamics of local neural signals, BEN is ideally suited to fill this specific void (Mauro & Wang, 2025; D.-H. Song & Z. Wang, 2024; Song, 2024; D. Song & Z. Wang, 2024). A few studies have started to document the developmental features of BEN, from age-related changes in neonates (Zhao et al., 2024) to comprehensive lifespan patterns from 8 to 89 years (Del Mauro, Zeng, & Wang, 2025). However, the current research landscape clearly lacks coverage of the vital developmental window between infancy and 8 years of age. Our work seeks to address this omission. By utilizing movie-watching fMRI data from participants aged 3 to 12, this study aims to bridge a portion of this critical gap and characterize the early developmental trajectory of BEN.

## 2. Methods

### 2.1 Participants

All data utilized in the study were obtained from the publicly available OpenNeuro database, specifically dataset ds000228 (https://openneuro.org/datasets/ds000228/versions/1.1.1), which was released by Richardson et al (2023). This database includes 122 children (M±SD = 6.7±2.3, 64 females) aged 3.5 to 12 years and 33 adults (ages 18–39 years; M±SD = 24.8±5.3,20 females) participants. Written consent was obtained from all adult participants, and both parental/guardian consent and child assent were secured for all child participants. The recruitment and experimental procedures were approved by the Committee on the Use of Humans as Experimental Subjects (COUHES) at the Massachusetts Institute of Technology (MIT).

### 2.2 MRI acquisition

Whole-brain structural and functional MRI were acquired on a 3-Tesla Siemens Tim Trio scanner at the Athinoula A. Martinos Imaging Center at MIT, with custom head coils employed for children under age 5. Structural MRI data were collected via a T1-weighted sequence (176 interleaved sagittal slices; 1mm isotropic voxels). Functional MRI data were obtained with a gradient-echo EPI sequence (32 slices; TR=2s; TE=30ms; flip angle=90°), employing prospective acquisition correction for motion, and a total of 168 volumes were acquired. Minor variations in functional voxel size and slice gap existed due to the participants’ diverse recruitment origins. Consequently, all functional data were upsampled to 2mm isotropic voxels in normalized space for consistency.

Participants watched a silent version of ‘Partly Cloudy,’ a 5.6-minute animated movie, during functional MRI scanning. A short description about the film can be found online (https://www.pixar.com/partly-cloudy#partly-cloudy-1). Before this, participants aged five and older had completed additional tasks that largely entailed listening to stories (children) or reading stories (adults). The stimulus itself was then presented after a 10-second rest period, during which all participants were instructed to watch the movie and remain still.

### 2.3 MRI preprocessing

FMRI data were analyzed utilizing SPM8 in conjunction with custom code imple mented in Matlab and R by Richardson et al (2018). The spatial preprocessing pipeline involved a sequence of registrations: functional images were initially realigned to th e first image of the run, which was subsequently coregistered to the participant’s high-resolution anatomical image. After this, each anatomical scan was normalized to the Montreal Neurological Institute (MNI) template.

The fidelity of each individual’s brain registration to the MNI template was rigorously assessed via visual inspection, ensuring a precise match of the cortical outline and key internal neuroanatomical structures, including the AC-PC axis and major sulcal patterns. A final spatial smoothing step was applied to all functional volumes using a Gaussian filter with a 5 mm full-width at half-maximum (FWHM) kernel.

For more detailed participants’ information, MRI acquisition and preprocessing can be found in the original article from the dataset (Richardson, Lisandrelli, Riobueno-Naylor, & Saxe, 2018).

### 2.4 BEN mapping

The voxel-wise BEN maps were calculated from the preprocessed rs-fMRI images using the BEN mapping toolbox (BENtbx) (Wang et al., 2014) based on sample entropy (SampEn) (Richman & Moorman, 2000). SampEn is a measure designed to quantify irregularity of a time series by assessing the probability of similar patterns occurring within the sequence. It evaluates the likelihood that two sequences of length *m* and *m*+1 will remain similar under a given tolerance threshold *r*. In the study, the window length (dimension) was set to *m* = 3, and the cut-off threshold was set to *r* = 0.6 according to Wang et al (2014) experimental optimization after evaluating the impact of a range of window lengths (*m*) and threshold values (*r*) on the results (Wang et al., 2014). The first four volumes were discarded for signal stability. BEN maps were smoothed with an isotropic Gaussian kernel with FWHM□=□8 mm. More details of the BEN calculation can be found in the original BEN paper (Wang et al., 2014) or in other previous studies (Lin, Chang, Song, Li, & Wang, 2022; Liu et al., 2020; Song et al., 2019; Wang, 2021; Wang & Initiative, 2020).

### 2.5 Statistical analysis

To mitigate potential artifacts arising from the uneven absence of data in edge signals, such as those in the cerebellum and parietal cortex, all data were first multiplied and then binarized. A gray mask was subsequently applied and used as the unified mask for all subsequent analyses. The mean global BEN (gBEN) value was extracted from this mask. Inter-group differences between adults and children were assessed using a two-sample t-test. Furthermore, the correlation between age and the mean BEN was evaluated separately for each group. For these analyses, custom Python scripts were used, and statistical significance was defined as p < 0.05.

The voxel-wise inter-group differences in BEN between adults and children, along with the voxel-wise BEN-age correlations within each group, were performed by Statistical Parametric Mapping (SPM12, Wellcome Trust Centre for Neuroimaging, London, UK, http://www.fil.ion.ucl.ac.uk/spm/software/spm12/) (Friston et al., 1994). To assess relative changes in BEN, the values were also first standardized using z-scores and then scaled to a range of 0 to 1. For these analyses, statistical significance was defined using a voxel-level threshold of *p* < 0.005 combined with a cluster-size threshold of 20 voxels.

## 3. Results

### 3.1 Mean Global BEN

The mean gBEN in children is significantly lower than that in adults (*t* = −3.228, *p* = 0.002, Cohen’s *d* = −0.561), even after controlling for sex and handedness (*t* = −2.944, *p* = 0.004). However, no significant correlation was found between age and mean gBEN in either children (*r* = −0.057, *p* = 0.534, *n* = 122) or adults (*r* = 0.130, *p* = 0.473, *n* = 33), even after accounting for sex and handedness (children: *r* = −0.0417, *p* = 0.6485, adults: *r* = 0.1233, *p* = 0.4942). Figure 1 presents the distribution of mean gBEN for each participant and its relationship with age.

**Figure 1.**
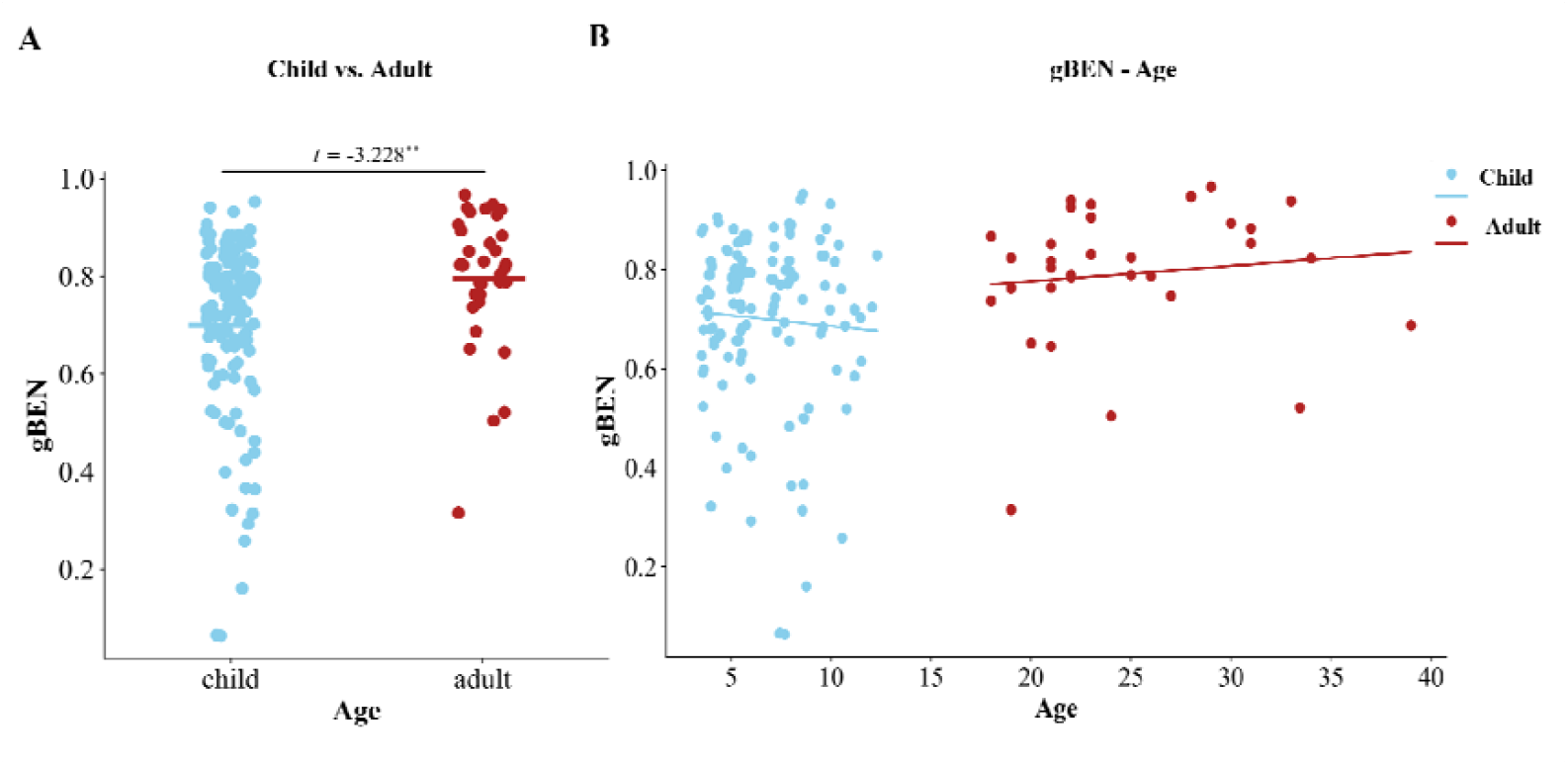
Age-related differences and correlations in mean global BEN. (A) Differences in mean global BEN between children and adults. (B) Scatter plot showing the distribution of mean global BEN across different age groups, with the fitted trend lines. Data points for children and adults are colored blue and red, respectively.

### 3.2 Voxel-wise BEN

Compared to adults, children exhibited lower BEN in several regions, including the orbitofrontal cortex (OFC), posterior cingulate cortex (PCC), fusiform gyrus (FFG), and thalamus (Fig.2A). Furthermore, in children, BEN in the right insula (INS), dorsomedial prefrontal cortex (DMPFC), and anterior cingulate cortex (ACC) were negatively correlated with age, a relationship not observed in adults (Fig.2B). Mean BEN values extracted from these three regions all demonstrated significant negative correlations with age in children (INS: *r* = −0.294, *p* < 0.001; DMPFC: *r* = −0.293, *p* = 0.001; ACC: *r* = −0.275, *p* = 0.002), even after controlling for gender and handedness (INS: *r* = −0.286, *p* = 0.001; DMPFC: *r* = −0.279, *p* = 0.002; ACC: *r* = −0.269, *p* = 0.003). In contrast, no significant associations between age and BEN were observed in these regions in adults (INS: *r* = 0.095, *p* = 0.597; DMPFC: *r* = 0.244, *p* = 0.172; ACC: *r* = 0.3000, *p* = 0.090), even after accounting for the same covariates (INS: *r* = 0.116, *p* = 0.522; DMPFC: *r* = 0.255, *p* = 0.152; ACC: *r* = 0.283, *p* = 0.110). Figure 2C presents the distribution of mean BEN and the corresponding fitted curves for both children and adults in these regions.

**Figure 2.**
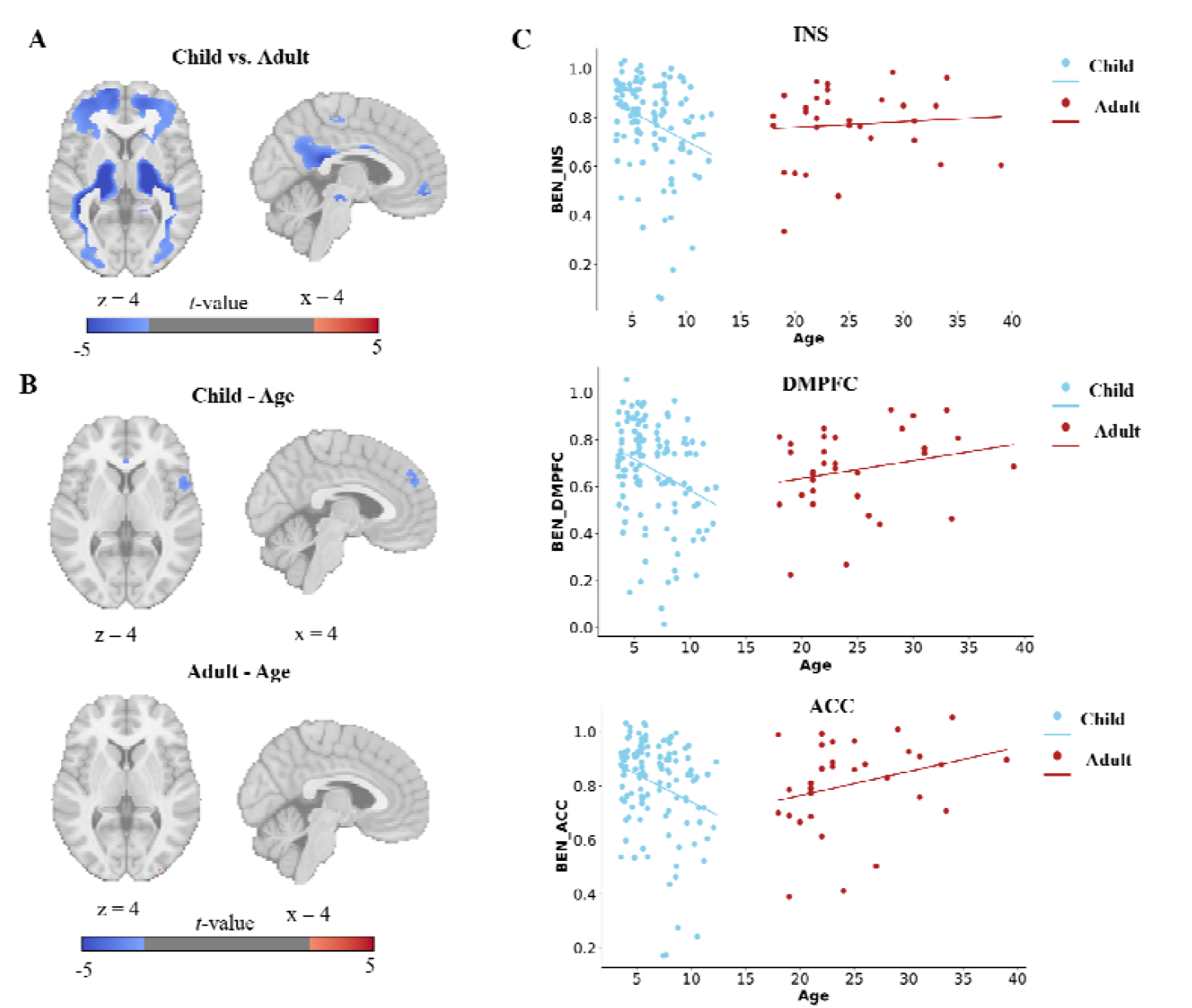
Age-related differences and correlations in voxel-wise BEN. (A) Differences in voxel-wise BEN between children and adults. Blue indicates lower BEN in children. (B) Correlations of voxel-wise BEN-age for children and adults, respectively. Blue indicates negative correlation. (C) Scatter plot showing the distribution of mean voxel-wise BEN from INS, DMPFC, and ACC across different age groups, with the fitted trend lines. Data points for children and adults are colored blue and red, respectively.

### 3.3 Standardized BEN

To examine the relative changes in BEN across different brain regions, we spatially standardized the BEN maps for each participant. The relative gBEN is higher in children than in adults, and relative gBEN is negative with age in children (*r* = −0.252, *p* = 0.005). The results of voxel-wise analysis revealed that children had higher relative BEN than adults in most brain regions, including the medial prefrontal cortex (MPFC), cingulate cortex (CC), sensorimotor cortex (SMC), temporal cortex (TC), visual cortex (VC), and striatum (Fig.3A). Furthermore, in children, relative BEN showed significant negative correlations with age in the INS, DMPFC, ACC, striatum, thalamus, and ventral tegmental area (VTA) (Fig.3B). In contrast, adults exhibited a significant positive correlation in the right dorsolateral prefrontal cortex (DLPFC) and a negative correlation in the precuneus (PCu) (Fig.3B). Subsequent analysis of the INS, DMPFC, and ACC confirmed that relative BEN demonstrated stronger negative correlations with age than absolute BEN in these regions (INS: *r* = −0.499, *p* < 0.001; DMPFC: *r* = −0.473, *p* < 0.001; ACC: *r* =−0.447, *p* < 0.001), even after controlling for gender and handedness (INS: *r* = −0.498, *p* < 0.001; DMPFC: *r* = −0.465, *p* < 0.001; ACC: *r* = −0.451, *p* < 0.001). In contrast, no significant associations between age and BEN were observed in these regions in adults (INS: *r* = 0.085, *p* = 0.639; DMPFC: *r* = 0.292, *p* = 0.099; ACC: *r* = 0.336, *p* = 0.056), even after accounting for the same covariates (INS: *r* = 0.083, *p* = 0.648; DMPFC: *r* = 0.277, *p* = 0.118; ACC: *r* = 0.294, *p* = 0.097). Figure 3C presents the distribution of mean standardized BEN and the corresponding fitted curves for both children and adults in these regions.

**Figure 3.**
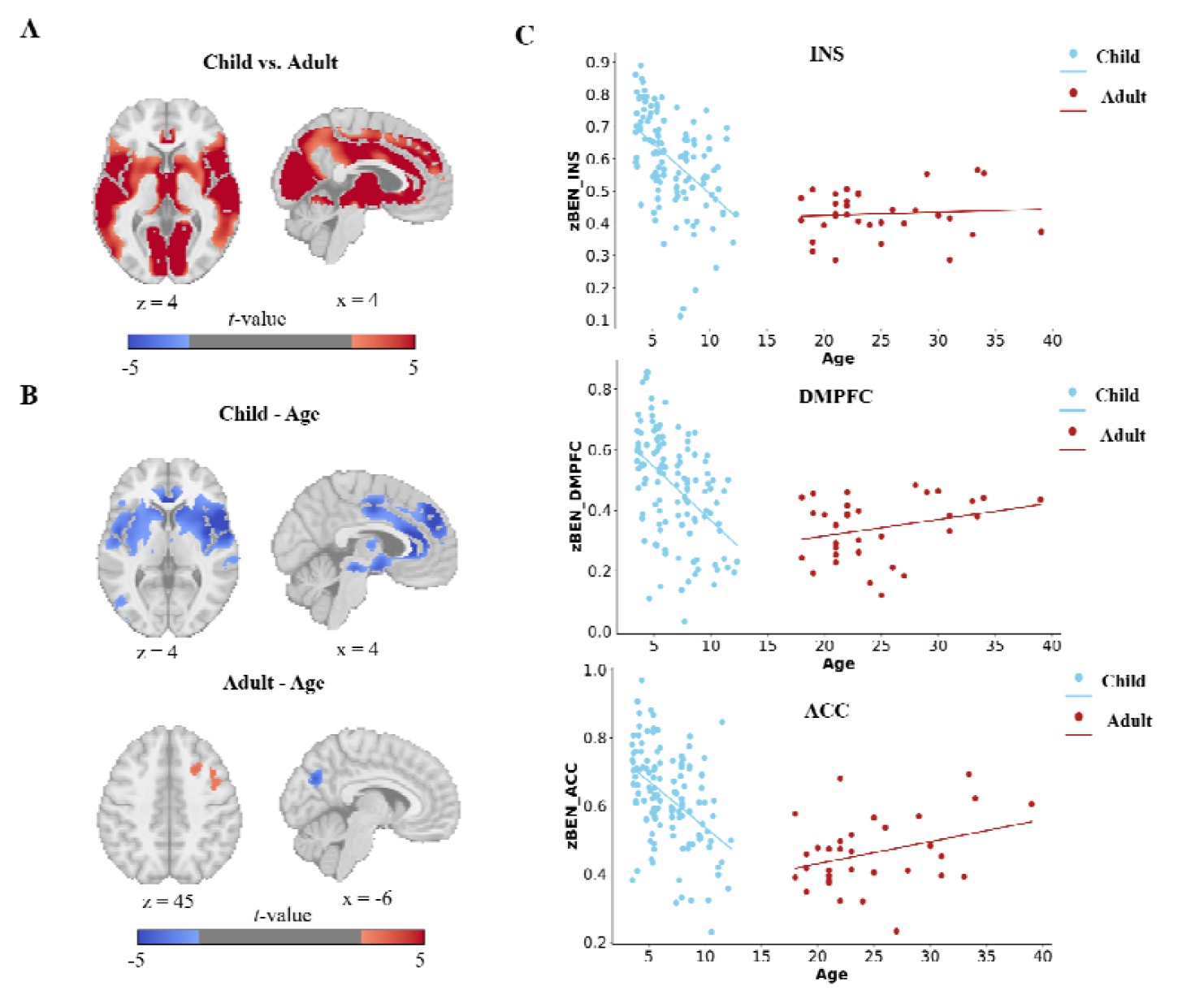
Age-related differences and correlations in standardized BEN. (A) Differences in standardized BEN between children and adults. Red indicates higher BEN in children. (B) Correlations of standardized BEN-age for children and adults, respectively. Blue indicates negative correlation, red indicates positive correlation. (C) Scatter plot showing the distribution of mean standardized BEN from INS, DMPFC, and ACC across different age groups, with the fitted trend lines. Data points for children and adults are colored blue and red, respectively.

## 4. Discussion

The preliminary findings of this study indicate that children aged 3-12 years exhibit lower gBEN compared to adults. Interestingly, no significant correlation was found between age and whole-brain BEN within either group. Voxel-wise analysis revealed lower BEN in children within several regions, primarily including the OFC, PCC, FFG, and thalamus. Subsequently, a significant negative correlation between age and voxel-wise BEN was identified in children, specifically in the INS, DMPFC, and ACC, a pattern not observed in adults. Spatially normalized BEN analysis revealed that children had significantly higher relative BEN in multiple brain areas, including the MPFC, CC, TC, VC, and striatum. In children, relative BEN was negatively correlated with age in the INS, DMPFC, ACC, striatum, thalamus, and VTA. In adults, relative BEN showed a positive correlation with age in the right DLPFC and a negative correlation in the PCu.

Regarding gBEN, lower values in children suggest that their overall brain activity is more coherent and predictable. This indicates a fundamentally simpler and less differentiated global neural architecture compared to the highly complex and specialized adult brain. Voxel-wise analysis localized these significant reductions in BEN primarily to key nodes of the default mode network (DMN) (Davey, Pujol, & Harrison, 2016; Menon, 2023), such as the medial OFC and PCC. The DMN is critically involved in internal thought processes, including self-reflection, autobiographical memory, future prospection, and social cognition, particularly theory of mind. Consequently, the lower BEN observed in these regions may be functionally linked to the relative immaturity of these advanced social-cognitive abilities in children.

In contrast, the analysis of relative BEN (spatially normalized) revealed a more dynamic picture. Children exhibited higher relative BEN in widespread regions, including large portions of the sensorimotor and visual cortices. This elevated relative entropy in primary processing areas may reflect the intense, exploration-driven interaction children have with their environment. The external world, being novel and rich with learning opportunities, demands heightened neural processing in these regions, making their activity patterns relatively less predictable as they engage with and learn from sensory inputs.

Furthermore, within the child group, a significant negative correlation between age and BEN was identified in the INS, DMPFC, and ACC, which are core hubs of the action-mode network (AMN)(Badke D’Andrea et al., 2025; Dosenbach, Raichle, & Gordon, 2025). This negative correlation was even stronger for relative BEN. The AMN supports goal-directed behaviors, requiring heightened arousal, focused attention, and the planning/execution of actions. The age-related decrease in BEN within this network may signify the progressive refinement and internalization of cognitive control. As children develop, their goal-directed behaviors become more efficient and automated, leading to more predictable and streamlined neural activity within the AMN.

Collectively, these findings demonstrate that relative BEN is a particularly sensitive marker for capturing developmental changes in brain function. It is highly plausible that these functional entropy dynamics are underpinned by profound structural maturation. Recent neuroimaging studies, such as those from the brain charts for the human lifespan, indicate that cortical grey matter volume peaks around age 5.9 years (Bethlehem et al., 2022), followed by a prolonged period of synaptic pruning and refinement. Therefore, future analyses would be significantly strengthened by incorporating structural metrics such as grey matter volume to directly investigate the influence of brain morphology on the developmental trajectory of BEN.

In summary, this preliminary study robustly demonstrates that BEN during movie-watching is a sensitive and informative biomarker, capable of reflecting key characteristics of functional brain development in children aged 3-12 years.

## Acknowledgements

We thank the Athinoula A. Martinos Imaging Center at the McGovern Institute for Brain Research at MIT, Jorie Koster-Hale, Natalia Velez-Alicea, Mika Asaba, and Nir Jacoby for help with data collection. We thank Todd Thompson for his work in making this dataset publicly available.

## Data and code availability

The original data is available in the online repository of OpenNeuro ds000228 (https://openneuro.org/datasets/ds000228/versions/1.1.1).

The BENtbx is available at https://www.cfn.upenn.edu/zewang/BENtbx.php and https://github.com/zewangnew/BENtbx.

